# TFAP2A+ embryonic progenitor cells undergo fate diversification to give rise to human amnion, germline, and mesoderm

**DOI:** 10.1101/2025.06.26.661839

**Authors:** Auriana Arabpour, Jonathan Adam DiRusso, Qiu Ya Wu, Mark Larsen, Young Sun Hwang, Elsie Jacobson, Thi Xuan Ai Pham, Nicole Agranonik, Megan Sparrow, Vernon Leander Monteiro, Zenya Rebecca Bian, Nicolas Pelaez-Restrepo, Antuca Callejas-Marin, Vincent Pasque, Kathrin Plath, Amander T. Clark

## Abstract

Amnion, germline and mesoderm specification at the posterior end of the human embryo occur around the same time *in vivo*. Similarly, *in vitro* generation of germline and amnion is associated with mesoderm induction regardless of differentiation platform. Yet, the lineage relationships between amnion, germline and mesoderm remains unresolved. By adding Basement Membrane Extract (BME) to the media, we demonstrate emergence of TFAP2A+/SOX2-epithelial progenitor cells which develop in response to BMP receptor signaling. We track the order of embryonic events that take place from this progenitor pool revealing that amnion-like cells (AMLCs) and primordial germ cell (PGC)-like cells (PGCLCs) are specified first. Shortly after, gastrulating mesoderm-like cells (MeLCs) arise that undergo an epithelial to mesenchymal transition (EMT). These results highlight the interconnected role of basement membrane deposition and BMP receptor signaling in the specification of human germline, amnion and mesoderm from TFAP2A+ embryonic progenitors.

## Introduction

The second and third week of human embryogenesis, during which implantation and lineage specification occurs, is critical to life. Abnormalities at these stages can result in pregnancy loss, infertility and birth defects. The Carnegie collection, which uses morphological features to stage human embryo development^1^, provided some of the earliest insights into this window of human life. These studies described a bilaminar disc consisting of two epithelial layers: the epiblast atop the hypoblast. During implantation, a proamniotic cavity develops from the epiblast leading to specification of extraembryonic amniotic cells and PGCs^1–3^. Concurrently, the primitive streak (PS) forms at the most posterior end of the epiblast disc, thus marking the start of gastrulation.

Beyond morphological features of rare human embryos, our current understanding of this developmental window is pieced together from eutherian model organisms^4^, human pluripotent stem cell (PSC) differentiation, human stem cell-based embryo models^5,6^, human embryo attachment cultures^7^, single cell profiling^8^ and spatial transcriptomics of human samples^9,10^.

Comparative studies of amnion formation indicate that this window of development post-implantation has diverged significantly amongst eutherians^11,12^. For instance, in human and non-human primates (NHPs), the amniotic cavity is created through a cavitation process in tandem with amnion induction, preceding gastrulation. However, in other eutherian embryos such as the mouse, rabbit and porcine, amnion induction occurs through a folding mechanism during gastrulation^13^. Such developmental divergence in peri-implantation embryogenesis may have functional consequences for the molecular events involved in early embryonic lineage specification around the time of amnion formation.

One such example is the origin and segregation of PGCs which occurs around the time of embryo implantation. Given this is a challenging window of human development to study, other mammalian embryos including mouse, NHP, rabbit and porcine are used as alternate models, with each species revealing unique insights into PGC development. For example in the mouse embryo, where the epiblast develops as an egg cylinder, PGCs develop from BLIMP1+ PGC-precursors in the posterior epiblast, giving rise to PRDM14+/PRDM1+/TFAP2C+ specified PGCs in the allantoic mesoderm^1^. In rabbit embryos which develop as bilaminar discs, OCT4+/TFAP2C+ putative PGCs are first identified in the posterior pre-streak epiblast before localization to the emerging mesoderm upon gastrulation^14^. In other studies, PGCs were first identified in the hypoblast of primitive-streak stage rabbit embryos^15^. In bilaminar disc porcine embryos^16^, SOX17+ PGCs are first identified in the posterior pre-streak epiblast^17^ consistent with the one report in the rabbit. Evaluating old-world (cyno) and new world (marmoset) NHP bilaminar disc embryos reveals a slightly different origin. In both primate species, specified PGCs were first observed in the amnion at embryonic day 11/CS5, and then at the boundary between amnion and posterior embryonic disc at CS6^18,19^. Transcriptional profiling of specified cyno, marmoset and human PGCs reveals conserved gene expression patterns across these species^20^. However, functional studies targeting the ape-specific^21^ long terminal repeat 5 Human Specific (LTR5Hs) transposon sequences suggests a new mechanism has evolved in human PGC induction compared to monkeys^22^. Given these similarities and differences, understanding human PGC specification benefits from incorporating human stem cell-based models.

Modeling human PGC development *in vitro* involves the induction of PGCLCs from human PSCs. The original cytokine and chemical-driven models were designed for high-yield induction of PGCLCs in 3D disorganized aggregates^23,24^. More recently monolayer and 2D induction protocols were developed for this same purpose^25,26^. The creation of geometrically confined 2D micropatterned colonies from hPSCs were amongst the first in vitro models to reveal the close spatial relationship between amnion, PS-like/mesoderm, and PGCLC induction upon BMP4 stimulation^27–29^. Similarly, 3D stem cell-based embryo models have also pointed to a spatial relationship between specified PGCs, pre-streak epiblast, amnion and hypoblast ^30–33^. At a molecular level, comprehensive single-cell transcriptomic datasets of *in vitro* human PGCLC induction^34^, and diffusion pseudotime ordering of marmoset embryo development^20^ independently revealed TFAP2A+ embryonic progenitor cells as likely precursor cells of the human germline. These embryonic progenitor cells develop in response to BMP signaling, and preceding expression of the PGCLC/PGC transcription factors SOX17 and TFAP2C^20,34^. Null mutations in *TFAP2A* identified a requirement for TFAP2A in PGCLC induction and a modest effect on mesoderm formation but no significant effect on amnion cell formation^20^. However, using an amnion-specific induction protocol, TFAP2A was found to control subsequent progression of amniogenesis^35^. Given that TFAP2A effects PS/mesoderm differentiation, it is also possible that some mesodermal cells likely develop from a TFAP2A+ embryonic progenitor pool as well.

In recent work, we demonstrated that simply adding BME to hPSC media as an overlay two hours after single cell plating was sufficient to promote amnion, germline and mesoderm cell fate specification in a model called Gel-3D^36^. Here, we report a 2D BME overlay model that uses chemically defined Essential 8 (E8) medium to study upstream modifiers of PGCLC induction, and the downstream lineages that emerge from TFAP2A+ embryonic progenitors (Fig. 1a). Our model develops AMLCs, PGCLCs, PS-like cells (PSLCs), MeLCs and endoderm-like cells (EndLCs), with MeLCs being the predominant cell type by day 3. Given the low-density plating of this model, it is highly conducive to live cell imaging. This allows for visualization of the dynamic self-instructive process of embryonic cell differentiation in the absence of added cytokines beyond TGFb1/Nodal and FGF2 found in E8 together with BME overlay. We have named this new model Geltrex-incorporated Medium Overlay (GiMO).

**Fig. 1:**
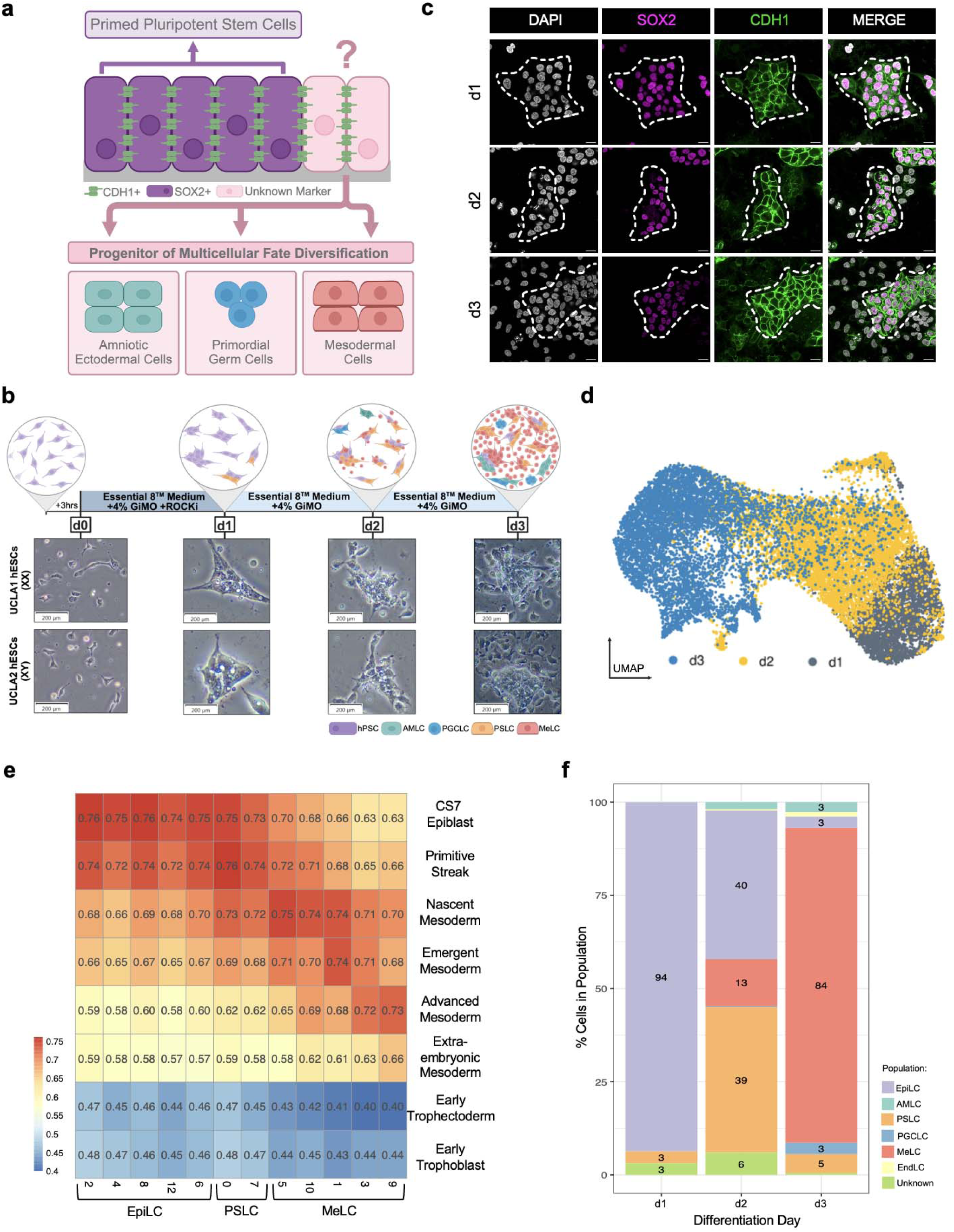
GiMO self-instructing differentiation recapitulates the peri-gastrulating epiblast. **a,** A schematic of GiMO differentiation. Schematic was made using BioRender.com. **b,** A schematic (top) and bright field representative images (bottom) of GiMO differentiation per day in UCLA1 (XX) and UCLA2 (XY) hESCs. Scale bars = 200 μm. Schematic was made using BioRender.com. **c,** Representative images of GiMO-generated colonies at d1-d3. SOX2 (magenta) marks epiblast cells, CDH1 (green) marks E-cadherin positive colonies, and DAPI (gray) marks all nuclei present. Scale bar = 20 μm. SOX2, CDH1, and DAPI had their intensities increased for clarity. **d,** UMAP plot of scRNA-seq data reveals 3 clusters, each corresponding to the day of GiMO differentiation (d1-d3). **e,** heatmap of correlation coefficients from integrated scRNA-seq datasets from pre-implantation, post-implantation and CS7 human embryo and specified distinct clusters generated from d1-3 GiMO. **f,** Relative proportions of cell populations identified by distinctly generated clusters per day of GiMO differentiation

## Results

### GiMO gives rise to cells that are specified in the posterior end of the human embryo

GiMO begins with human PSCs cultured in E8 media on vitronectin coated plates. Following digestion to single cells, the PSCs are seeded at a density of 12,500 cells cm^-2^ on a 1% coat of Geltrex in E8 media (Fig. 1b). To stimulate lineage differentiation, the media is changed 3 hours (hrs) after seeding to E8 media containing 4% Geltrex. This media is replaced at 24hrs (day 1; d1) and 48hrs later (d2) with experiments ending at d3 or a maximum of 72hrs post plating. At d1, the PSCs form colonies. By d2, single migratory cells can be observed leaving the colonies, the number of which increase greatly and becoming highly migratory between d2 and d3 (Fig. 1b, Supplementary Video 1).

Based on what morphologically appears to be the formation of epithelial colonies in the first 24 hrs of GiMO, we next performed immunofluorescence (IF) to characterize expression of the epithelial protein E-cadherin (CDH1) together with the epiblast marker SOX2 (Fig. 1c). This data shows that at d1 of GiMO, cells have formed CDH1+ colonies composed of SOX2+ cells. By d2 and d3, the cells leaving the colonies are negative for CDH1 as well as SOX2 suggesting lineage differentiation is coincident with EMT. To determine whether differentiation in GiMO is due to the addition of Geltrex to the media or plating cells at low density, we repeated the experiments with and without the 4% Geltrex overlay. We find that, in the absence of overlay, cells reform as PSCs (Extended Data Fig.1a-b). This result demonstrates that addition of 4% Geltrex to the media is necessary to trigger EMT in the GiMO model.

To further evaluate the dynamics of CDH1+ colony morphogenesis and EMT we used time-lapse microscopy focusing on the d1-d2 transition (Extended Data Fig. 1c). This data reveals that EMT begins at ∼44 hrs post overlay consistent with the timing of CDH1 repression by d2 GiMO. In addition, we also noticed that many of the colonies develop a lumen-like structure starting just before EMT at ∼32hrs post overlay (Extended Data Fig. 1c). To confirm whether the colonies retain lumens, we stained for EZRIN together with CDH1 at d3. We found that in CDH1+ colonies almost 80% exhibit EZRIN in the center (Extended Data Fig. 1d-e). Taken together, we propose that the GiMO model creates CDH1+ EpiLC colonies that undergo lumenogenesis and EMT; all events that occur in early peri-implantation epiblast morphogenesis ^37–40^.

Next, to examine the identity of cells generated in GiMO, we performed single-cell RNA-sequencing (scRNA-seq) using 10X Genomics at d1, d2 and d3. After performing integration and Uniform Manifold Approximation and Projection (UMAP) across all three days we identified 16 clusters representing progressive changes in cell identity from d1-d3 (Fig. 1d and Extended Data Fig. 2a). Using diagnostic genes, we assigned embryonic lineage classifications to those clusters which included epiblast-like cells (EpiLCs), PSLCs, AMLCs, PGCLCs, MeLCs and EndLCs (Extended Data Fig. 2a-b). To further characterize the 16 clusters based on *in vivo* embryonic development, we integrated the 10X data with existing single-cell datasets from preimplantation, post-implantation and a Carnegie Stage 7 (CS7) human embryo datasets^8,41–43^ (Extended Data Fig. 2c). Using this approach, cells generated in GiMO are highly comparable with epiblast, amnion, PGCs, PS, primitive endoderm (PE) and mesoderm but not to trophoblast cells (Fig. 1e, Extended Data Fig. 2b and Extended Data Fig. 2d-e). Based on this, we were able to reduce the 16 clusters into six distinct embryonic cell types (Extended Data Fig. 2f).

Subsequently, we were particularly interested in whether GiMO generates extraembryonic mesoderm-like cells (ExMLCs) as reported in prior ECM-overlay studies^26^. By comparing our MeLC population to previously defined ExM datasets^44^, we show that the mesoderm generated in GiMO is embryonic (Fig. 1e and Extended Data Fig. 2f). We also discovered a small population of “unknown” cells at d1 and d2 that express low levels of pluripotent genes (Extended Data Fig. 2a-b and Fig 1e). These cells do not correlate with any *in vivo* embryonic cell types and may represents an artefact of the *in vitro* system, or a rare post-implantation *in vivo* progenitor population that has not yet been identified.

Taken together, over the first three days of GiMO the following dynamic embryonic events are modelled (Fig 1f): d1, 94% of cells correspond to EpiLCs with an emerging population of PSLCs. By d2 PSLCs become more prominent (39%), and MeLC begin to emerge (13%). In addition, rare populations of AMLCs, PGCLCs and EndLCs are also first identified. By d3, the vast majority of GiMO cells (84%) corresponds to MeLCs. Additionally, as EpiLCs and PSLCs diminish from d2-d3 transition, PGCLCs, AMLCs, EndLCs are also present but are rapidly outcompeted by MeLCs. Thus, GiMO is a 2D model that largely recapitulates EpiLC morphogenesis and EMT giving rise to mostly MeLCs.

### GiMO captures gastrulation-like EMT and formation of mesoderm

In the developing CS7 embryo, three types of gastrulating mesoderm can be identified: nascent, emergent, and advanced^18^ (Fig 2a). Based on these previously characterized *in vivo* mesodermal markers, we can identify all three populations of MeLCs in GiMO (Fig. 2b). Nascent MeLC is an early PS mesoderm co-expressing *TBXT* (PS marker) and *MESP1* (mesoderm marker). Emergent MeLC lacks *CDH1* while expressing *LHX1* and *MESP1.* Advanced MeLC retains *LHX1*, expresses *HAND1* and *BMP4* as well as EMT markers *SNAI2, ZEB1, ZEB2 and MMP2*^8^. We find that nascent MeLCs (*CDH1+/TBXT+/GATA6+*) emerge by d2 GiMO while emergent and advanced MeLCs (*GATA6*+/*CDH*1-) emerge more prominently by d3.

**Fig. 2:**
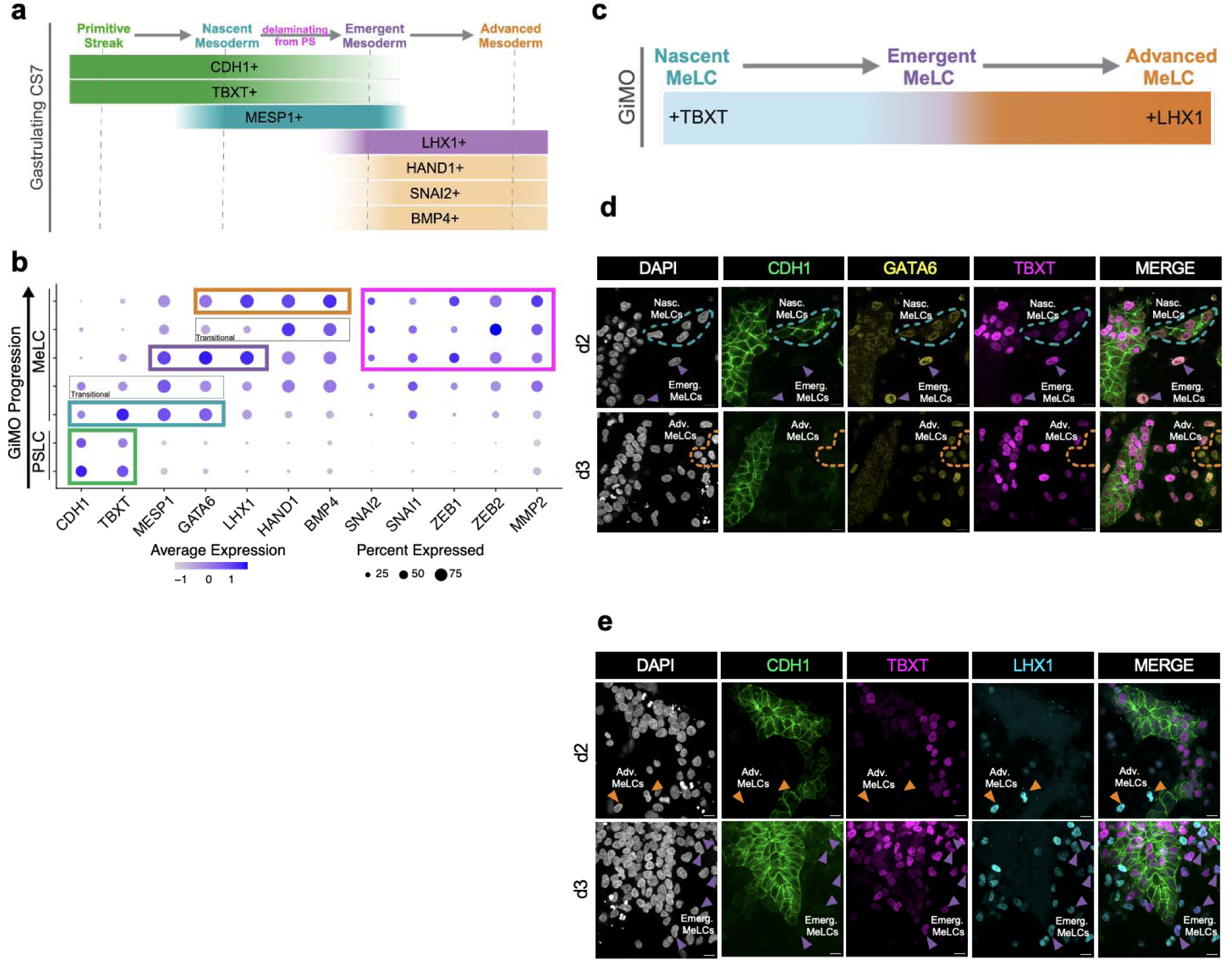
GiMO captures gastrulation-like EMT of mesoderm specification. **a,** A schematic of marker expression during CS7 gastrulation EMT. Schematic was made using BioRender.com. **b,** Dot plot analysis of key markers across scRNA-seq clusters associated with PSLC and MeLC populations across GiMO progression. **c,** A schematic of marker expression during GiMO gastrulation-like EMT. Schematic was made using BioRender.com. **d,** Representative images of gastrulating MeLCs d2 and d3 GiMO. CDH1 (green), GATA6 (yellow) and TBXT (magenta) mark nascent MeLCs while GATA6 and TBXT together negative for CDH1 mark emergent MeLCs. GATA6 only positive cells marks advanced MeLCs. Scale bar = 20 μm. CDH1, GATA6, TBXT, and DAPI (gray) had their intensities increased for clar ty. **e,** Representative images of gastrulating MeLCs d2 and d3 GiMO. LHX1 (cyan) together with TBXT (magenta) marks emergent MeLCs while LHX1 only positive cells marks advanced MeLCs. Scale bar = 20 μm. CDH1 (green), GATA6, TBXT, and DAPI (gray) had their intensities increased for clarity.

To spatially position nascent, emergent, and advanced MeLCs in GiMO, we stained for GATA6 together with CDH1, TBXT, and LHX1 (Fig. 2c-e). Aligning with our scRNA-seq data, we found that at d2, nascent MeLCs (CDH1/TBXT/GATA6 triple positive) are located primarily in the epithelial colonies. By d3, emergent and advanced MeLCs correspond to single cells outside the colonies (Fig. 2d). Cells with bright or dim TBXT+ expression together with bright or dim GATA6 or LHX1 expression likely represent the transitional populations of nascent to emergent MeLCs (Fig. 2d-e).

### TFAP2A induction and SOX2 repression are induced in response to BMPR signaling

Since GiMO recapitulates posterior cell types where BMP signaling is expected to be enriched and where BMP-responsive lineages first emerge^10^, we next investigated whether cells autonomously express *BMP4* in response to overlay. We were also interested in identifying cells that are likely competent to respond to BMP4 via expression of Type II BMP receptor (*BMPR2*). By projecting *BMP4* onto the UMAP, we found that *BMP4* is highly expressed by MeLCs at d3; after lineage differentiation has occurred (Extended Data Fig. 2g). To identify *BMP4* expressing cells at earlier time points, we analyzed cell populations at d1 and d2 of GiMO. We discovered that about one-quarter of EpiLCs at d1 express *BMP4,* and that at d2, *BMP4* is expressed by EpiLCs, PSLCs, AMLCs, and MeLCs (Fig. 3a). Next, to identify cell types that are BMP4-responsive across the GiMO time course, we examined *BMPR2* expression and found that most cells at d1-3 express *BMPR2* and are likely competent to respond to BMPs (Fig. 3b).

**Fig. 3:**
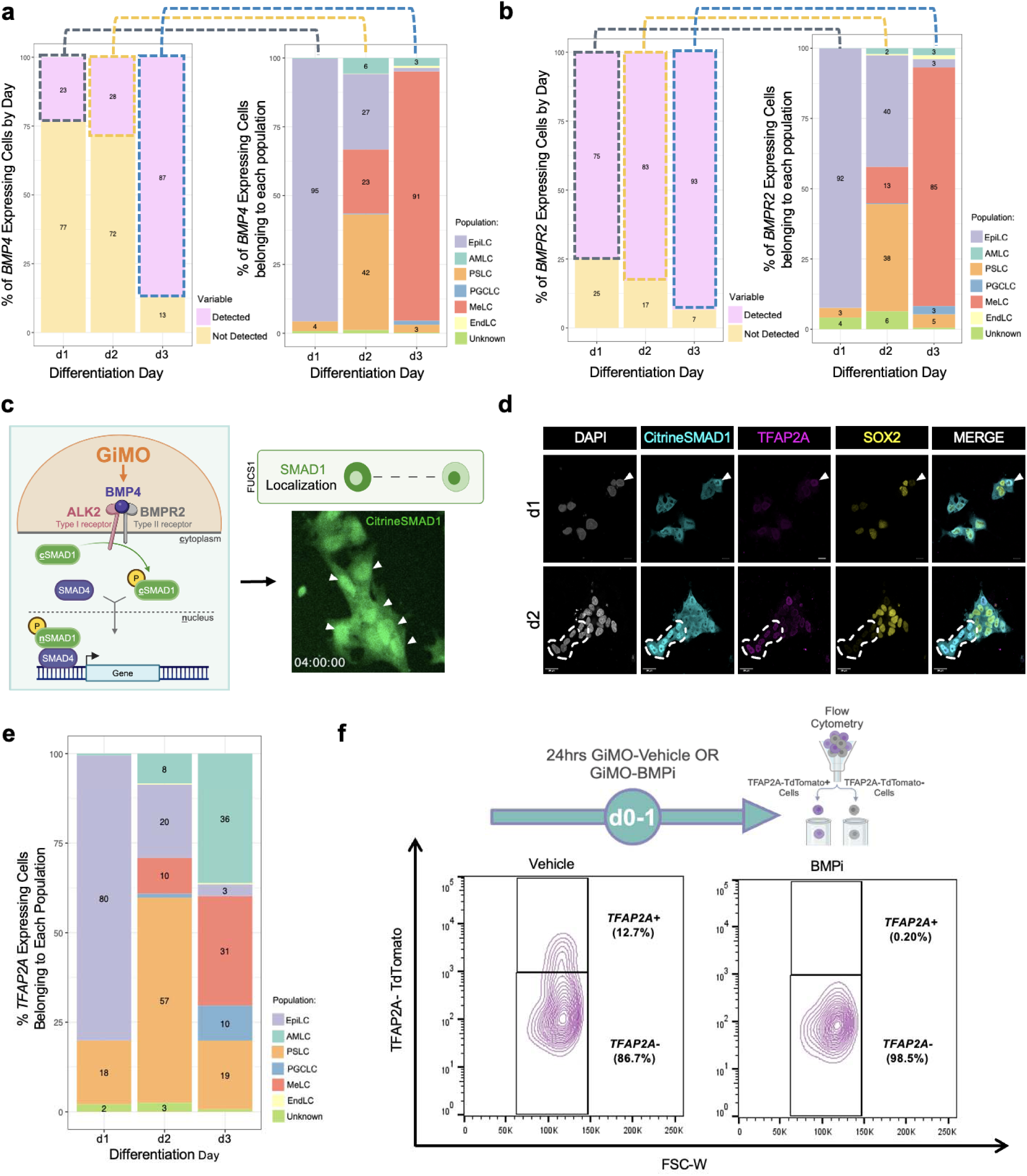
GiMO induces early SMAD1 nuclear localization and TFAP2A expression. **a,** The percentage of cells expressing *BMP4* by cluster (left) and relative proportions of cells belonging to each population (right) that’s detected to express *BMP4* per day of GiMO differentiation. **b,** The percentage of cells expressing *BMPR2* by cluster (left) and relative proportions of cells belonging to each population (right) that’s detected to express *BMPR2* per day of GiMO differentiation. **c,** A schematic of BMP4-responsiveness in GiMO (left) and representative image of SMAD1 nuclear localization using FUCS1 hiPSCs at d0 GiMO. Schematic was made using BioRender.com. **d,** Representative images of SMAD1 nuclear localization d1 and d2 GiMO. Citrine-SMAD1 (anti-GFP, cyan) and TFAP2A (magenta) together mark BMP4-responsive cells. Scale bar = 30 μm. Citrine-SMAD1, TFAP2A, SOX2 (yellow), and DAPI (gray) had their intensities increased for clarity. **e,** Relative proportions of cells belonging to each population that’s detected to express *TFAP2A* per day of GiMO differentiation. **f,** Experimental design (top) and flow cytometry analysis (bottom) at d1 GiMO of TFAP2A-TdTomato expressing cells from BMPi (5μm DMH1 concentration) or Vehicle (DMSO equivalent volume). Schematic (top) was made using BioRender.com.

To visualize BMP responsiveness in GiMO under live-cell conditions, we used a F-lentivirus hUbiC > Citrine-SMAD1 (FUCS1) human iPSC reporter line engineered to express Citrine tagged SMAD1 protein under the human Ubiquitin-C constitutive promoter. In the self-renewing state when human PSCs are cultured in E8 media on vitronectin, we found no nuclear localization of the fusion protein indicating that prior to GiMO, human PSCs in E8 media are not signaling via SMAD1 translocation into the nucleus (Extended Data Fig. 3a). In contrast, within 4hrs of overlay, we observed nuclear translocation of Citrine-SMAD1 (nSMAD1) in most cells within colonies visualized via live-cell conditions (Fig. 3c, Supplementary Video 2). This responsiveness is consistent with most cells expressing *BMPR2* and being competent to respond to BMP receptor signaling via nSMAD1. Interestingly, we also noticed that SMAD1 translocation was dynamic, with some cells within colonies retaining nSMAD1 while for others Citrine-SMAD1 was rapidly lost from the nucleus (Supplementary Video 2).

Next, to evaluate an early BMP-responsive embryonic gene, we performed IF for TFAP2A^35,45,46^ together with SOX2, given that SOX2 expression is often reduced following exposure to BMP4^14,17,47^. To identify cells with nSMAD1 we used an anti-GFP antibody to detect Citrine. Our results show that within colonies, when SMAD1 is localized to the nucleus, expression of TFAP2A is coincident with repression of SOX2 (Fig. 3d). Given the loss of SOX2 in TFAP2A+ cells, we evaluated a second pluripotent transcription factor, NANOG, together with CDH1. At d1, TFAP2A protein is localized to subpopulations of CDH1+ cells that are dim or negative for NANOG. By d2, the majority of TFAP2A+ cells are still CDH1+ and NANOG is repressed (Extended Data Fig. 3b). Taken together, this result indicates that expression of TFAP2A is associated with repression of SOX2 and NANOG protein in CDH1+ cells.

Given that TFAP2A is often used as a marker of amnion and trophoblast, we were interested in the identity of *TFAP2A+* cells in GiMO. Using our scRNA-Seq dataset, we found subpopulations of cells at any given day which express *TFAP2A* (Fig. 3e). At d1 GiMO, ∼80% EpiLCs and ∼18% PSLCs expressed *TFAP2A*. By d2, a transition occurred where ∼20% EpiLCs and ∼57% PSLCs expressed *TFAP2*A, along with emerging 8% AMLCs and 10% MeLCs. Finally, at d3, increasing proportions of AMLCs and MeLCs expressed *TFAP2A* along with ∼10% PGCLCs, 19% PSLCs, and ∼3% EpiLCs. Critically, this result indicates that *TFAP2A* expression is likely much broader in early human embryonic cells than simply amnion, trophoblast and PGCs. To confirm this, we evaluated *TFAP2A-*expressing cells in existing *in vivo* human embryo datasets and show that *TFAP2A* is also expressed in subpopulations of epiblast, PS, amnion and PE (Extended Data Fig. 3c).

To further evaluate the dynamics of TFAP2A in GiMO, we created a TFAP2A-P2A-TdTomato reporter tool in hPSCs (TFAP2A-TdTomato) (Extended Data Fig. 3d). To confirm specificity to TFAP2A-expressing cells, we performed IF for TdTomato (anti-RFP) and TFAP2A to show that TFAP2A/RFP overlap (Extended Data Fig. 3e). Using flow cytometry for TdTomato+ cells we also found dynamic changes in the proportion of TFAP2A-TdTomato+ cells across the time course (Extended Data Fig. 3f-g). Therefore, like *TFAP2A* RNA (Fig. 3e), TFAP2A protein is likewise expressed in multiple subpopulations of embryonic cell types across d1-3 GiMO. To visualize TFAP2A-TdTomato expression under live-cell conditions, we used spinning disk time-lapse microscopy and identified TdTomato+ cells as early as 4hrs following overlay. Consistent with our IF and flow cytometry results, we found that TFAP2A-TdTomato expression was not identified in every cell but instead was dynamically expressed in subpopulations of cells from 4-54hrs (Supplementary Video 3).

Although TFAP2A is considered a BMP-responsive gene, we wanted to confirm this result in GiMO through perturbation experiments to block endogenous BMP receptor (BMPR) signaling. Since BMPR2 preferentially binds ALK2 to form a heterodimeric complex^48^, we exposed cells to the ALK2-specific inhibitor, DMH1 (BMPi) and control cells to DMSO (vehicle). Flow cytometry for TdTomato+ expression was performed 24hrs post overlay. We found no induction of TFAP2A-TdTomato signal in the first 24hrs of GiMO when compared to control (Fig. 3f). Taken together, this result demonstrates that the initial induction of *TFAP2A* in EpiLCs downstream of an ECM overlay, involves BMPR signaling, and the outcome of this initial TFAP2A induction is repression of SOX2.

### TFAP2A+ progenitors develop into germline and amnion cells

As SOX2 repression is required for PGCLC induction^14,17^, we then evaluated TFAP2A expression in PGCLCs^20,26,34^. Using IF, we first confirmed the presence of NANOG+/SOX17+ PGCLCs and positioned them within CDH1+ colonies at d2 and d2.5 (Extended Data Fig. 4a). By d3, we found PGCLCs to have significantly reduced levels of CDH1 protein (Extended Data Fig. 4b). Across all time points, single-positive NANOG cells were also identified likely representing EpiLCs that have not yet initiated differentiation while DAPI+ only cells represented other somatic lineages (Extended Data Fig. 4a). Next, to determine whether induction of PGCLCs is also associated with nuclear translocation of SMAD1, we performed IF for Citrine-SMAD1 together with NANOG and SOX17 to mark PGCLCs using the FUCS1 reporter line (Fig 4a). We found that NANOG+/SOX17+ PGCLCs co-expressed nSMAD1. Therefore, the induction of PGCLCs within GiMO colonies is associated with cells receiving and responding to BMPR signaling.

**Fig. 4:**
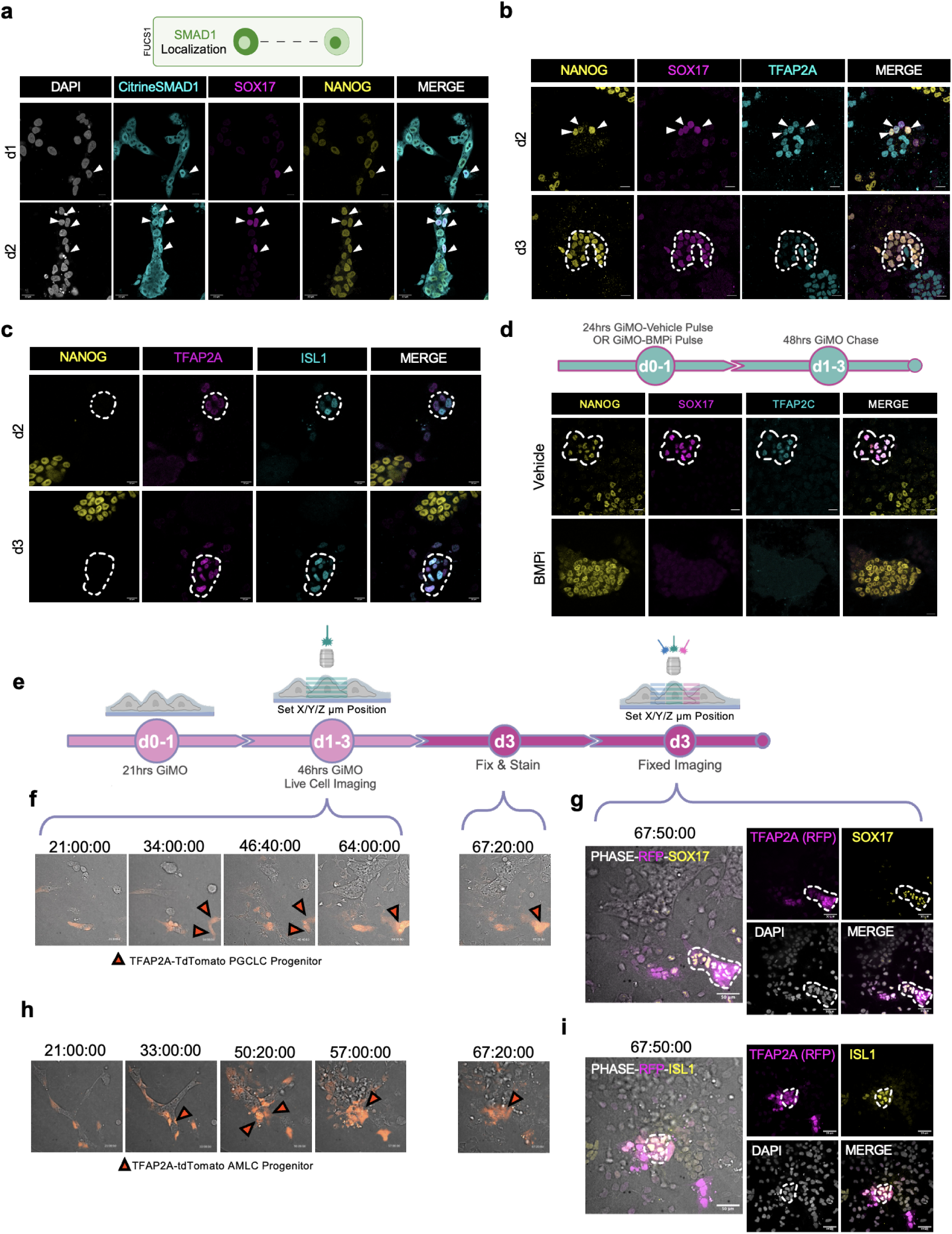
PGCLCs and AMLCs are induced from TFAP2A+ epithelialized cells. **a,** A schematic of FUCS1 hiPSC SMAD1 localization (top) and representative images (bottom) of SMAD1 nuclear localization d1 and d2 GiMO. Citrine-SMAD1 (anti-GFP, cyan) together with SOX17 (magenta) and NANOG (yellow) mark BMP4-responsive PGCLCs. Scale bar = 30 μm. Citrine-SMAD1, SOX17, NANOG, and DAPI (gray) had their intensities increased for clarity. Schematic (top) was made using BioRender.com. **b,** Representative images of PGCLCs expressing TFAP2A (cyan) at d2 and d3 GiMO. NANOG (yellow) together with SOX17 (magenta) mark PGCLCs. Scale bar = 50 μm. NANOG, SOX17, and TFAP2A had their intensities increased for clarity. **c,** Representative images of AMLCs expressing TFAP2A (magenta) at d2 and d3 GiMO. ISL1 (cyan) together with TFAP2A mark AMLCs. Scale bar = 20 μm. NANOG (yellow), TFAP2A, and ISL1 had their intensities increased for clarity. **d,** A schematic of the experimental design (top) and representative images (bottom) of PGCLCs at d3 GiMO under BMPi (5μm DMH1 concentration) or Vehicle (DMSO equivalent volume). NANOG (yellow) together with SOX17 (magenta) and TFAP2C (cyan) mark PGCLCs. Scale bar = 20 μm. NANOG, SOX17, and TFAP2C had their intensities increased for clarity. Schematic (top) was made using BioRender.com. **e,** A schematic of the live imaging and IF staining experimental design. Schematic was made using BioRender.com. **f,** Representative images of time-lapse videos showing reporter expression with phase contrast. TFAP2A (anti-RFP), SOX17 (yellow), phase contrast (gray) and DAPI (gray) had their intensities increased for clarity. **g,** IF stain from fixed live cell imaging of PGCLCs (yellow) together with TFAP2A-TdTomato progenitor (magenta). **h,** Representative images of time-lapse videos showing reporter expression with phase contrast. **i,** IF stain from fixed live cell imaging of AMLCs (yellow) together with TFAP2A-TdTomato progenitor (magenta). TFAP2A (anti-RFP), ISL1 (yellow), phase contrast (gray) and DAPI (gray) had their intensities increased for clarity.

Since TFAP2A is a downstream target of BMP signaling, we then evaluated TFAP2A protein expression in PGCLCs. Our data shows that at d2, NANOG+/SOX17+ PGCLCs co-express TFAP2A protein and by d3, TFAP2A protein expression is reduced (Fig. 4b). Given the close physical relationship between amnion and the initial wave of PGC induction in primates^18^, we evaluated TFAP2A together with the amnion marker ISL1 (Fig. 4c). This result demonstrates that like PGCLCs, TFAP2A+/ISL1+ AMLCs are identified by d2 of GiMO.

To confirm that PGCLC induction is dependent on BMPR signaling, we established the same experiment as Fig. 3f where BMPi was added for the first 24hrs of overlay. Following BMPi treatment, fresh media containing 4% Geltrex overlay was then added at d1 and d2 to create a pulse-chase experiment. By phase contrast microscopy, we showed that control cells undergo EMT (Extended Data Fig. 4c) and produce PGCLCs (Fig. 4d). In contrast, cells treated with BMPi for the first 24hrs remained as colonies and retained NANOG expression (Fig. 4d) with limited evidence of EMT (Extended Data Fig. 4c) or TFAP2A induction (Extended Data Fig. 4d). This result suggests that during the first 24hrs of GiMO, BMPR responsiveness is critical to NANOG repression, TFAP2A expression, PGCLC induction and EMT.

Given that PGCLCs and AMLCs originate from TFAP2A+ progenitors, we performed live-cell imaging with the TFAP2A-TdTomato reporter line followed by IF for PGCLC and AMLC markers. In this experimental design, spinning disk time-lapse microscopy was used focusing on ∼21hrs to ∼67hrs post overlay. Once live imaging was completed at pre-determined X/Y/Zμm positions, wells were fixed with 4% PFA, stained for PGCLC and AMLC markers, and re-imaged at the original X/Y/Zμm positions (Fig. 4e). This result reveals that PGCLCs emerged from a TFAP2A-TdTomato+ progenitor pool ∼34hrs post overlay (Fig. 4f-g, Supplementary Videos 4-5). Similarly, AMLCs also emerge from a TFAP2A-TdTomato+ progenitor ∼33hrs post overlay (Fig. 4h-i, Supplementary Videos 6-7). Given that PGCLCs and AMLCs have heterogenous origins (i.e. emerged from different EpiLC colonies), our results support the idea that PGCLCs and AMLCs are induced from a heterogenous TFAP2A+ progenitor pool and not a single cell with two fates.

### TFAP2A+ progenitors develop into highly migratory GATA6+ mesodermal cells

The third major cell type to form in GiMO are MeLCs, with most cells at d3 corresponding to GATA6+ MeLCs (Fig 1f, Extended Data Fig. 2b and Extended Data Fig. 2e). To first position GATA6+ cells in GiMO, we stained for GATA6 together with CDH1 and SOX2 (Fig. 5a). We found that GATA6+ MeLCs are first identified at d2 both within CDH1+ colonies, as well as in CDH1 negative cells. We also show that GATA6+ cells at all time points are negative for SOX2 (Fig. 5a).

**Fig. 5:**
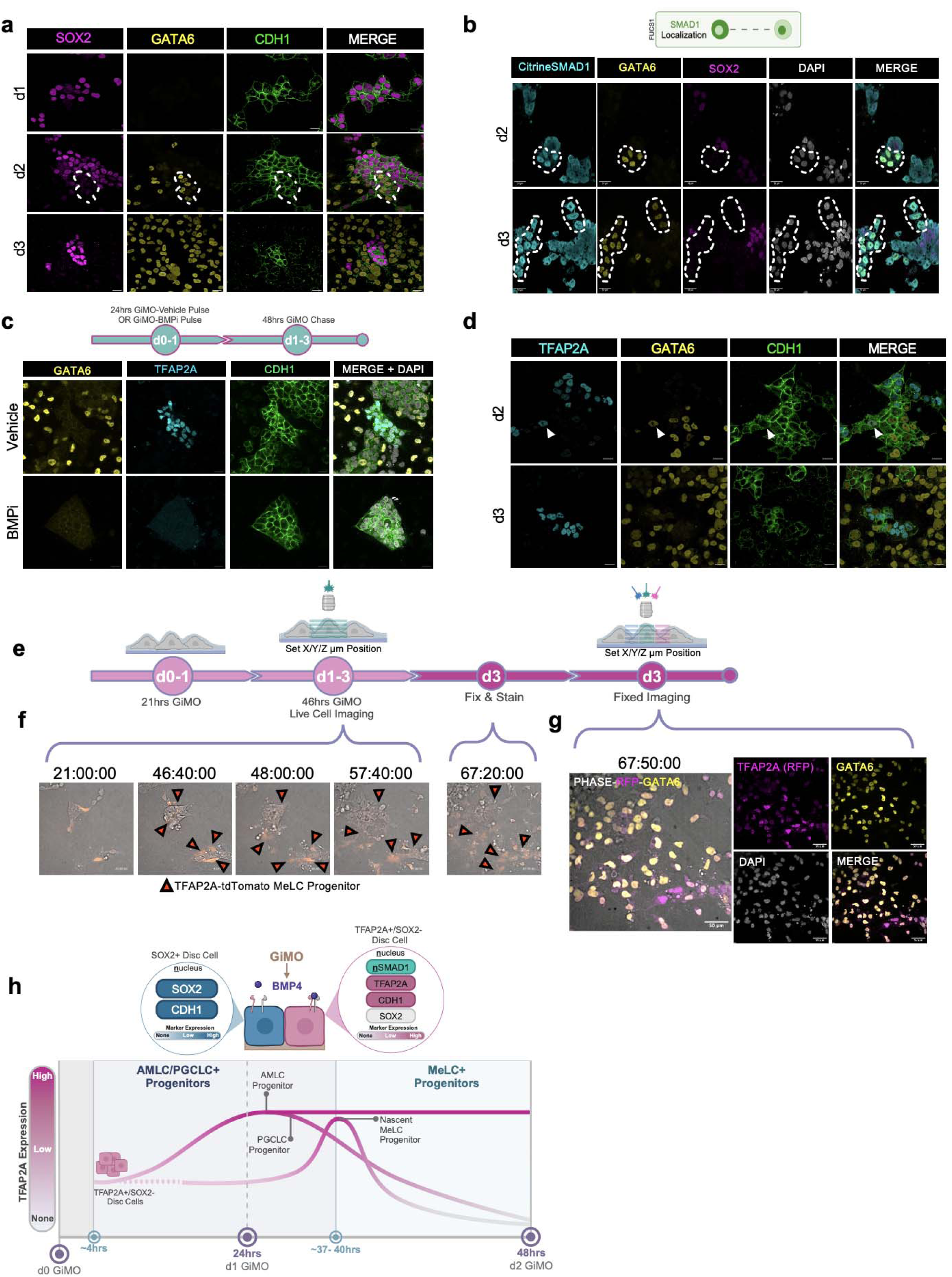
MeLC induction depends on BMPR signaling and form in TFAP2A+ epithelial colonies. **a,** Representative images of MeLCs d1 and d2 GiMO. GATA6 (yellow) marks MeLCs, SOX2 (magenta) marks EpiLCs, and CDH1 (green) marks colony cells. Scale bar = 50 μm. SOX2, GATA6, CDH1, and DAPI (gray) had their intensities increased for clarity. **b,** A schematic of FUCS1 hiPSC SMAD1 localization (top) and representative images (bottom) of SMAD1 nuclear localization d1 and d2 GiMO. Citrine-SMAD1 (anti-GFP, cyan) together with GATA6 (yellow) mark BMP4-responsive PGCLCs. Scale bar = 30 μm. Citrine-SMAD1, GATA6, NANOG (yellow), and DAPI (gray) had their intensities increased for clarity. Schematic (top) was made using BioRender.com. **c,** A schematic of the experimental design (top) and representative images (bottom) of MeLCs and TFAP2A+ cells at d3 GiMO under BMPi (5μm DMH1 concentration) or Vehicle (DMSO equivalent volume). Scale bar = 20 μm. GATA6 (yellow), TFAP2A (cyan), and CDH1 (green) had their intensities increased for clarity. Schematic (top) was made using BioRender.com. **d,** Representative images of GATA6+ and TFAP2A+ cells within colonies at d2 and d3 GiMO. Scale bar = 50 μm. TFAP2A (cyan), GATA6 (yellow), and CDH1 (green) had their intensities increased for clarity. **e,** A schematic of the live imaging and IF staining experimental design. Schematic was made using BioRender.com. **f,** Representative images of time-lapse videos showing reporter expression with phase contrast. TFAP2A (anti-RFP), GATA6 (yellow), phase contrast (gray) and DAPI (gray) had their intensities increased for clarity. **g,** IF stain from fixed live cell imaging of MeLCs (yellow) together with TFAP2A-TdTomato progenitor (magenta). **h,** A schematic of asynchronous emergence of BMP4-responsive progenitor cells that repress SOX2 and express TFAP2A. Schematic was made using BioRender.com.

Given that MeLCs are formed around the time of EMT, we next asked whether the EMT event was necessary for MeLC induction. To achieve this, we exposed cells to either an inhibitor of matrix metalloproteinases MMP2/9 (MMPi) or DMSO as a vehicle control (Extended Data Fig. 4e). In the vehicle control, GATA6+/CDH1-MeLCs were observed at d3 and were distinct from the CDH1+ colonies that express TFAP2A. In contrast, exposure of MMPi for three days resulted in GATA6+/CDH1+ MeLCs that were TFAP2A negative. Given that TFAP2A+ cells were identified in MMPi conditions, we also looked for PGCLCs and found that EMT is not required for induction of PGCLCs (Extended Data Fig. 4f).

Next, to evaluate whether induction of GATA6 is associated with BMPR signaling, we evaluated Citrine-SMAD1 nuclear localization by IF using the anti-GFP antibody together with GATA6 and SOX2 (Fig. 5b). This result shows that within colonies, the emergence of GATA6+ MeLCs is associated with SMAD1 nuclear localization, and in all cases, GATA6 expression is associated with SOX2 repression. Therefore, generation of GATA6+ MeLCs in colonies is likely a direct result of BMPR stimulation. To further confirm this, we performed pulse-chase experiments using the same format used previously (Fig 4d). In control conditions, most GATA6+ cells can be identified outside of CDH1+ colonies and are TFAP2A negative at d3. In BMPi pulse-chase conditions, only CDH1+ colonies are identified, with no expression of TFAP2A or GATA6 at d3 (Fig. 5c).

BMP stimulation is necessary for the induction of both TFAP2A+ and GATA6+ cells, yet GATA6+ cells are negative for TFAP2A. Therefore, to evaluate whether GATA6+ MeLCs express TFAP2A at an earlier time point, we stained GiMO at d2 and identified rare GATA6+ cells which are also positive for TFAP2A in CDH1+ colonies. However, the majority of GATA6+ MeLCs are TFAP2A negative (Fig 5d). To identify the precursors before GATA6 is induced, we performed live cell imaging with the TFAP2A-TdTomato reporter line using the same experimental design as before (Fig. 5e). We found that the earliest GATA6+ MeLCs emerge from TFAP2A+ cells ∼47hrs post overlay and that most of these cells repress TFAP2A once they leave the colony and begin rapidly migrating (Fig. 5sf-g, Supplementary Videos 8-9). After this first wave, however, migratory cells that ultimately express GATA6 did not originate from cells that express TFAP2A (Supplementary Movie 9). Thus, our data supports the hypothesis that a subpopulation of highly migratory GATA6+ MeLCs originate from TFAP2A+ cells, and that GATA6+ mesoderm can also be specified from TFAP2A negative cells as a second wave phenomena.

In sum, although GiMO does not involve the addition of BMP4 to the system, our results show that addition of overlay is sufficient to trigger embryonic cell differentiation which is dependent on BMPR signaling in the first 24hrs. This differentiation begins with asynchronous emergence of progenitor cells that repress SOX2 and express TFAP2A and are chronologically competent to first form AMLCs and PGCLCs. Shortly after, TFAP2A+/SOX2-progenitor colony cells become competent to then form the earliest GATA6+ MeLCs which are highly migratory (Fig. 5h).

## Discussion

Here, we generated a distinctly simple 2D model, GiMO, that reliably recapitulates posteriorized lineage diversification in the presence of FGF2 and TGFβ/Nodal (found in E8) together with a BME overlay. As this self-instructing model is conducive to imaging, the timing of early human lineage specification can be defined. We demonstrate that the emergence of TFAP2A+/SOX2-cells occurs at ∼4hrs post overlay, with induction of TFAP2A+ progenitors in the first 24hrs being reliant on BMPR signaling in the presence of FGF2/TGFβ/Nodal and BME. This leads to AMLC, PGCLC, and MeLC specification from d2-3 (Fig. 5h). We further observe that EpiLC colonies generate lumens at ∼32hrs and that at ∼33-34hrs AMLC and PGCLC progenitors begin to emerge. Next, initiation of EMT can be identified at ∼37-40hrs with generation of emergent/advanced MeLCs at ∼46hrs. After this point, CDH1-MeLCs rapidly fill the plate and become the major BMP4 producing cells of GiMO.

In our work, we show through single cell analyses and protein expression the link between BMPR signaling and SOX2 repression involves the induction of TFAP2A. Like the findings reported here and by others^20,26^, TFAP2A likely dampens the epiblast state to capacitate cells in the posterior embryo to engage in either amnion, PGC, or PS-nascent mesoderm fate decisions. Therefore, as the developing epiblast is exposed to a BMP-rich embryonic niche^10^, TFAP2A expression followed by SOX2 repression marks cells that will make these earliest embryonic lineage decisions. Furthermore, though we found CDH1+/SOX2+ cells and emerging CDH1+/TFAP2A+ cells to be mutually exclusive, we did capture rare NANOG+/TFAP2A+ colony cells that we hypothesize are cells committed to the PGCLC trajectory. In contrast to SOX17 and its role in PGCLC induction^24,49^, NANOG expression in the transition from epiblast to PGC progenitor remains to be resolved. Recent reports suggest that *NANOG* is continuously expressed in the posterior Epi(LC) to PGC(LC)- committed cells in both nonhuman primate embryos^18^ and in vitro derived human PGCLCs^25^. However, whether NANOG protein is transiently downregulated in human PGC (TFAP2A+) progenitors, requires further investigation.

Through our time-lapse experiments we find that mesodermal cells can originate from either a TFAP2A positive or TFAP2A negative progenitor that we hypothesize refer to the first and second wave of mesoderm induction^50^. We show via single cell analyses and protein expression that nascent MeLCs first emerge as BMP4-responsive cells within epithelialized colonies. As CDH1 negative MeLCs migrate, they become the dominant *BMP4*-expressing population that likely trigger TBXT+/TFAP2A-cells to induce MeLCs as a second wave phenomenon. Additionally, we find that blocking EMT by a MMP2/9 inhibitor does not impede induction of TFAP2A+ cells or formation of amnion, mesoderm or germ cells. This finding suggests that early embryonic lineage specification is independent to or precedes delaminate from the epiblast disc. Importantly, though EMT and ExM induction in the embryo are linked^44,51^, we find that the initiation and progression of gastrulation-like EMT does not rely on the presence of ExMLCs nor does it generate this type of mesoderm. Furthermore, although the exact origin(s) of ExM cells *in vivo* remains to be determined^44,52,53^, recent embryo models show that advanced MeLCs share some key characteristics with ExM cells^8,10,26,54^. Perhaps a subset of advanced mesodermal cells undergoing EMT in the 3D niche of embryogenesis acquire the capacity to also give rise to the ExM.

In summary we demonstrate that GiMO, a 2D self-instructing model, provides a reliable platform with which the molecular mechanisms driving lineage specification during peri-implantation can be dissected. Through GiMO, we’ve learned that BMPR signaling (nSMAD1+) is associated with SOX2 repression within epithelial colonies and expression of TFAP2A+ embryonic progenitors. These progenitors then go on to form AMLCs and PGCLCs followed by MeLCs that undergo gastrulation-like EMT. To our knowledge, our findings is the first to track the emergence of AMLC, PGCLC, and migratory MeLC using both a cytokine naïve medium and the lack of overexpression of TFs (via inducible reporters). We believe this to be a prelude in deciphering the developmental trajectories of cell populations differentiating in the early post-implantation human embryo.

## Methods

### Ethics Statement

This project uses de-identified human cell lines, specifically hESC and hiPSC lines. The hESC lines, called UCLA1 and UCLA2 were derived at UCLA from embryos consented for research following full committee review by the UCLA Medical Institutional Review Board (IRB) from the UCLA Office for Protection of Research Subjects. This committee reviewed the informed consent process, the consent form and all relevant documents involved in donation of surplus embryos to research at UCLA. The embryos were originally intended for family building purposes and were generated at an in vitro fertilization (IVF) laboratory until the donors chose to donate surplus embryos to research. Participants were not paid for participation, and all embryo donors were informed that the embryos would be destroyed as part of the research study. The derivation of hESC lines from human embryos was reviewed and approved by the UCLA IRB as well as the UCLA Embryonic Stem Cell Research Oversight Committee (ESCRO). UCLA1 and UCLA2 hESC lines were derived from blastocysts between D5-7 post-fertilization. The ESCRO committee did not request evidence of primitive streak formation during the hESC derivation process. The de-identified UCLA1 and UCLA2 hESC lines were provided to our lab from the UCLA human pluripotent stem cell bank following review and approval of the research by the UCLA human pluripotent stem cell research oversight committee (hpSCRO) (formally called ESCRO). The genetically modified hiPSC line called NPR00051 was received from the Elowitz lab at the California Institute for Technology following material transfer agreement (MTA). The parental hiPSC line used to generate NPR00051 is called WTC-11. This cell line was generated at UCSF before being deposited de-identified with the National Institute for Genome and Medical Sciences Human Genetic Cell Repository and distributed through Coriell Institute for Medical Research (Cell Line 25256). Experiments involving self-renewal and differentiation of hESC and hiPSC lines were reviewed and approved annually by the UCLA hpSCRO committee. All research with human embryos, hESC and hiPSC lines were performed following the principles laid out by the International Society for Stem Cell Research in the 2021 Guidelines for Stem Cell Research and Clinical Translation. This includes collaboration and communication with the UCLA hpSCRO committee who monitored the work, including specific endpoints for the experiments which included the minimal time necessary to achieve the scientific objective (in this case 3 days of differentiation).

### Cell Lines

hPSC lines used in this study include UCLA1 and UCLA2 human embryonic stem cells (hESCs)^55^, TFAP2A-TdTomato UCLA2 hESC, and FUCS1 hiPSC. All protocols for using these hPSC lines have been approved and reviewed annually by the UCLA Institutional Review Board (IRB #18-001466) together with annual review by the UCLA Human Pluripotent Stem Cell Research and Oversight (hPSCRO) Committee (hPSCRO #2018-005-04). All hPSCs were maintained in standard feeder-free culture for at least 17 passages and authenticated as karyotypically normal at the indicated passage number. The hPSC lines used in this study were all under P70 and all lines tested negative for mycoplasma contamination (MycoAlert Detection Kit, Lonza, # LT07-318).

### Cell Culture

hPSCs were maintained in standard feeder-free conditions using E8 medium (ThermoFisher Scientific, # A1517001) and cultured on vitronectin coated (VTN-N, ThermoFisher Scientific, # A14700) 6-well tissue culture plates (Corning, # 3516). hPSCs were passaged as aggregates every 6-7 days using an enzyme-free reagent for dissociation and passaging, ReLeSR (Fisher Scientific, # NC2331323). Cells were first washed with dulbecco’s phosphate-buffered Saline (D-PBS; Invitrogen, # 14190144) and 1mL of ReLeSR added to wells at room temperature (RT) for 1min. Next, 1mL of ReLeSR was aspirated and tissue culture plate placed in 37 °C and 5% CO_2_ incubator for an additional 4mins. After incubation, E8 Media was added (media volume based on desired passage dilution) per well to lift and dissociate hPSCs into aggregates and passaged onto freshly coated vitronectin plates.

### Generation of GiMO Culture

hPSCs used for GiMO experiments were first maintained until ∼65-70% confluency in 5-6 days of culture. Once ready, cells were washed with D-PBS and incubated with a 1:1 mixture of TrypLE Select enzyme (Gibco, # 50-591-419) and 0.5 mM EDTA/PBS solution (Nacalai USA, pH 8.0, # 13567-84) (0.5X TrypLE Select) at 37 °C, 5% CO2 for 10 min. Cells were quenched with E8 medium containing 10 μM ROCK inhibitor (Stemolecule Y27632, Fisher Scientific, # NC0577320) and gently pipetted to dissociate to single cells. Cell suspension was centrifuged at 1,500 rpm for 5 min before resuspending resultant cell pellets gently in E8 medium containing 10 μM ROCK inhibitor. Resuspended cells were then further dissociated to single cells using EASYstrainer Cell Sieves 40 μm filter (Greiner bio-one, VWR # 89508-342). Cell counts of all hPSCs were done manually using 4-chip disposable hemocytometers (Bulldog Bio Products, # DHC-N420) and counted ∼3 times per line to average total cell counts for high accuracy. After counts were completed, EASYstrainer Cell Sieves 40 μm filter was used again to ensure single cell plating. Each hPSC line was then plated in each well at 12,500 cells cm^-2^ in technical triplicate in E8 medium containing 10 μM ROCK inhibitor. Prior to cell plating, 24-well tissue culture plates (Corning, # 3526) were pre-coated with 1% Lactate dehydrogenase-elevating virus (LDEV)-free, hESC-qualified reduced growth factor basement membrane matrix Geltrex (ThermoFisher Scientific, # A1413302) for at least 1.5hrs at 37 °C and 5% CO_2_ based on manufacturer’s recommendations for hPSC culture plate coating. After seeding, plates were incubated for 3-3.5hrs at 37 °C and 5% CO_2_. Finally, culture medium (E8 medium containing 10 μM ROCK inhibitor) was changed to fresh E8 medium containing 4% Geltrex and 10 μM ROCK inhibitor. Culture medium was replenished daily thereafter. ROCK inhibitor was removed from culture medium at 24hrs after initial cell seeding.

### Generation of TFAP2A-TdTomato hESC Reporter Line

UCLA2 hESCs were cultured on plates coated with Recombinant laminin-511 E8 (iMatrix-511 Silk, Amsbio, # 892021) and were maintained under feeder-free conditions in StemFit Basic03 medium (Ajinomoto, # SFB-503) containing basic FGF (Peprotech, # 100-18B) at 37 °C and 5% CO_2_. Prior to passaging, hESC cultures were treated with 0.5X TrypLE Select for 15 min at 37 °C and 5% CO_2_ to dissociate into single cells. For routine maintenance, hESCs were plated into a 6-well plate at a density of 200,000 cells cm^-2^ with 10 µM ROCK inhibitor (Y-27632; Tocris, # 1254) added in culture medium for 1 day after hESCs passaging.

To construct the donor vector for generating the *TFAP2A-p2A-tdTomato* allele, homology arms of *TFAP2A* (left arm: 1480 bp; right arm: 916 bp) were first PCR-amplified from the genomic DNA of UCLA2 hESCs and were sub-cloned into pCR-Blunt II-TOPO vector using Zero Blunt TOPO PCR Cloning Kit (Thermo Fisher Scientific, # K280020SC). A p2A-tdTomato sequence and a PGK-Neo antibiotic resistance cassette flanked by loxP sites were inserted at the 3’-ends of TFAP2A homology arms using the NEBuilder HiFi DNA Assembly kit (New England Biolabs, # E2621S). Oligonucleotide duplexes corresponding to the following sgRNA sequences targeting PAM sites around the 3’-end of TFAP2A’s coding region (CGCCAAAAGCAGTGACAAAG and TTTTGGCGTTGTTGTCCGTG) were designed by the Molecular Biology CRISPR design tool (Benchling) and cloned into pX335-U6-Chimeric BB-CBh-hSpCas9n (D10A) SpCas9n-expressing vector to generate the sgRNAs/Cas9n vector (Addgene, # 42335)^56^.

The TFAP2A-p2A-tdTomato donor vector (2.5 ug) and sgRNAs/nCas9 vectors (1.25 ug each) were introduced into 800,000 UCLA2 wild-type hESCs by nucleofection using a P3 Primary Cell 4D-Nucleofector X Kit L (Lonza, # V4XP-3024). Single colonies were isolated after selection with neomycin (150 ug/mL) and Monoallelic UCLA2 TFAP2A-p2A-tdTomato reporter lines were selected for subsequent cell line establishment. Neomycin-resistant cells were then transfected with 2.5 ug of a plasmid containing a Cre-recombinase encoding sequence to excise *PGK-Neo* cassette. The success of the targeting, random integration and Cre recombination process was assessed by performing PCR on cell lysates from each colony. All the reporter lines used within this study have been characterized as karyotypically normal analyzed by G-banding performed by Cell Line Genetics (Madison, WI). Reporter hESC lines were gradually adapted to the E8/Vitronectin culturing system before GiMO differentiation.

### Generation of FUCS1 hiPSC Reporter Line

The human SMAD1 protein was fluorescently tagged by inserting a monomeric Citrine N-terminal tag on SMAD1. The Citrine-SMAD1 fusion protein was cloned into a F-U-lentiviral vector backbone, with a human Ubiquitin-C constitutive promoter^57^. DNA fragments were cloned using the Gibson Assembly Cloning Kit (NEB, # E5510S) in accordance with the manufacturer’s instructions. The resulting DNA was subsequently used to package the FUCS1 lentiviral plasmid to create functional lentiviral particles in HEK293FT cells using Delta and VSV-G plasmids^57^. Viral titer of the resulting concentrated particles was assessed through serial dilution in HEK293T cells.

Viral particles carrying the lentiviral FUCS1 DNA were transduced into WTC-11 hiPSCs (Coriell cell line 25256) cultured in mTeSR1 (Stem Cell Technologies, # 85850) supplemented with Penicillin-Streptomycin (Sigma-Aldrich, # P4333) using Matrigel (Corning, # 354277) as a substrate. Lentiviral transduction was carried by plating approximately 1×10^5^ WTC-11cells with 10µM Y-27632 (Stem Cell Technologies, # 73302). Cells were passaged using ReLSR (Stem Cell Technologies, # 100-0483) prior to generation of a clonal line for analysis. The clonal line was produced by dissociating cells in 100µl Trypsin-EDTA (0.05%) and re-replated sparsely (∼30-50 cells/well) in Matrigel coated 6-well plates preincubated with mTeSR1 supplemented with CloneR (Stem Cell Technologies, # 05888). Colonies were allowed to grow for 5-7 days before lifting individual clones manually with a P-10 pipette under a microscope and transferring each colony to individual wells of a 24-well plate.

### Immunofluorescence

hPSCs were fixed in 4% paraformaldehyde (PFA, Thermo Fisher Scientific, # 28908); buffered in 1X D-PBS at RT for 1hr and permeabilized in 0.1% sodium dodecyl sulfate (SDS, Fisher Scientific, #AAJ60015AC) for 40 min on a shaker. hiPSC lines were then blocked in 10% normal donkey serum (NDS, Sigma-Aldrich, # S30-100ML) for 1.5-2hrs at RT, followed by incubation with primary antibody solutions in 10% NDS at 4 °C on a shaker for 12 - 16hrs. Samples were then labeled with donkey-raised secondary antibodies (1:250 dilution) in 4% NDS at RT for 45 min. Cell nuclei were stained with 4,6-diamino-2-phenylindole (DAPI; ThermoFisher Scientific, # D1306). All primary antibodies, their sources, and dilutions are listed in Supplementary Table 1. All experiments that required IF and confocal imaging were done in 24-well black frame plates with high performance #1.5 cover glass (0.170±0.005mm) (Cellvis, # P24-1.5H-N).

### Quantification of Immunofluorescent Images

Analysis of CDH1 fluorescence intensity was performed using FIJI (ImageJ) v2.14. Briefly, PGCLC and non-PGCLC clusters (as identified by SOX17/NANOG double positive) were determined and circled manually. From PGCLC+/PGCLC-selected ROIs the average intensity was taken. Within each image, the ratio of intensity between PGCLC+ and PGCLC-clusters was taken as the ratio to control for image-to image variability. Data was collected across three replicates and plotted using R 4.3.3/ggplot2 (3.5.1). Statistics by ggpubr 0.6.0 using method “t.test”.

### BMP and EMT Inhibition Assays

In the small-molecule inhibitor treatment assays, either 5 μM BMP inhibitor DorsoMorphin Homolog 1 (DMH1; Cayman Chemical, # 16679), or 30 μM MMP2/MMP9 inhibitor I (abcam, # ab145190) was added to the culture medium at indicated time points. Dimethylsulfoxide (DMSO; Sigma-Aldrich, # D2650) was added to the control groups.

### scRNA-seq and Data Analysis

GiMO culture at d1, 2 and 3 were first washed with cold D-PBS 3X in 30sec increments to dissolve Geltrex overlay. Next, samples were treated with 0.25% Trypsin/EDTA for 10 mins at 37 °C and 5% CO_2_. After incubation, cells were quenched with cold E8 medium and centrifuged at 1,200 rpm for 3.5 min. Cells were then resuspended with FACS buffer, D-PBS containing 0.5% Bovine Serum Albumin (BSA, Sigma-Aldrich, # A3311), as a wash step and centrifuged at 700 rpm for 4 min. Cells were resuspended again with FACS buffer and dissociated to single cells using a Falcon 100 μm Cell Strainer (Corning, # 352360) and manually counted. hPSCs were then further dissociated using EASYstrainer Cell Sieves 40 μm filter and centrifuged at 300 rpm for 3 min and resuspended in FACS buffer at a total volume of 1000 cells μL^-2^. Within 1hr after cell dissociation, cells were loaded into the 10x Genomics Chromium.

Gene expression matrices for each time point were generated with CellRanger (v5.0.1 – 10X genomics), using STAR^58^ to align reads to the hg38 reference genome. The count matrices were thresholded and analyzed with Seurat (v4.3.0)^59^. Low quality cells and doublets were excluded from analysis by filtering out cells with fewer than 2500 or greater than 10000 unique genes, as well as any cell with >20% mitochondrial reads. The time points were then merged and the merged dataset was normalized with NormalizeData, top 2000 most variable features identified with FindVariableFeatures, and scaled with ScaleData. PCA was run and principle components assessed with ElbowPlot and DimHeatmap, and the top 15 principle components were used for dimensionality reduction and visualization with UMAP. The top 15 principle components were used to construct a nearest neighbor graph with FindNeighbors, and clusters were identified with a resolution of 1 using FindClusters. Differentially expressed genes marking each cluster were identified with FindAllMarkers, and clusters were identified using known markers of each cell type.

For proportion analysis, the expression of each gene across all clusters was checked using RidgePlot and a cutoff chosen. We then found the proportion by taking the number expression/total cells for each day. We then subset the dataset using this cutoff. For total percentage expressing a gene of interest, the number of cells from the subsetted object for each day (d1-d3) was divided by the number of cells total d1-d3 from the non-subsetted object. Dotplots were made with the DotPlot using standard arguments.

For correlation coefficient between the CS7 PGCs, CS7 Yolk Sac Endoderm (ArrayExpress E-MTAB-9388), UCLA1 and UCLA2 convention aggregate PGCLCs (GSE GSE140021) and GiMO PGCLCs (this study), 10X genomics datasets (conventional aggregate and GiMO) were aligned using CellRanger v5.0.1 to the hg38 genome. The Tyser et al. CS7 dataset was aligned using STARSolo (v2.7.9a) using the same hg38 genome and calling arguments -- soloType SmartSeq, --soloStrand Unstranded, --outFilterScoreMin 30, --soloUMIdedup Exact, --outFilterMultimapNmax 500, --outSAMattributes NH HI AS nM, --winAnchorMultimapNmax 100, --outSAMmultNmax 1. For GiMO and conventional aggregates, NANOS3-expressing cells were taken as PGCLCs and read counts from these cells aggregated using Seurat AggregateExpression with return.seurat = F. Yolk Sac Endoderm and PGCs from Tyler et. Al. were identified as in (CITE LTR5Hs paper). Read matrices for each population were imported into Seurat using ReadSTARSolo and aggregated using AggregateExpression. Aggregated counts from all experiments were combined and all genes with no reads in any one condition removed. Count depth was normalized by EdgeR (v3.38.4) using calcNormFactors. CPM values were generated using EdgeR cpm, normalized.lib.sizes = T. Pearson correlation of the resulting CPM values was calculated and plotted using pheatmap (v1.0.12).

### Integrated Analysis of Multiple Datasets

Single-cell RNA-seq datasets^8,41–43^ were integrated with the newly generated data from this study using the standard Seurat v3 integration workflow^60,61^. Prior to integration, the datasets were normalized and scaled, followed by the selection of the 2,000 most variable genes. Integration anchors were identified using the FindIntegrationAnchors function with default parameters, ensuring consistent incorporation of features and data across all samples. A Uniform Manifold Approximation and Projection (UMAP) for integrated data visualization was constructed using the RunUMAP function, specifying dimensions 1 to 10. Additionally, the DoHeatmap function was employed to generate a heatmap for detailed visualization of expression patterns.

### Live Cell Video Acquisition

GiMO live cell imaging experiments were conducted using the Zeiss Confocal composed of a Yokogawa CSU X1 spinning disk confocal on an inverted Zeiss stand. Fluorescence images were recorded with a 10x objective. A GFP filter set was used for live fluorescent imaging of FUCS1 cells. For TFAP2A-TdTomato experiments a Phase and RFP filter set was used for live fluorescent imaging. The live fluorescent imaging ran an exposure time of 150 ms (FUCS1) or 100 ms (TFAP2A-TdTomato) and was performed on 20 pre-determined X/Y/Zμm positions across 3-wells of a chambered coverslip with 4-wells and a #1.5H glass bottom (Ibidi, # 80427) with 15 (FUCS1) or 25 (TFAP2A-TdTomato) Z stacks and an interval of 20 mins rest per position.

### Detection of PGCLCs, AMLCs and MeLCs After Live Cell Imaging

After live cell imaging of TFAP2A-TdTomato reporter line was completed, the samples were fixed, permeabilized, blocked, and stained either for anti-RFP, SOX17 and DAPI to detect PGCLCs; or anti-RFP, ISL1, and DAPI to detect AMLCs; or anti-RFP, GATA6 and DAPI to detect MeLCs in GiMO culture. Plate was re-imaged at pre-determined X/Y/Zμm positions using the same Zeiss Yokogawa CSU X1 spinning disk confocal with the following laser settings: 561 laser with emission filter 627/62, 639 laser with emission filter 690/50, and 405 laser with emission filter 450/50.

### Live Cell Video Analysis

Videos were processed using ImageJ/FIJI version 2.41. Briefly, z-stacks were merged and brightness was adjusted for each channel and scale bars and time stamps added. Videos were exported directly from FIJI using JPEG compression at 10 frames per second.

### Flow Cytometry

Cells cultured for specified days and experiments were first washed with cold 1X D-PBS three times in 30sec increments to dissolve Geltrex overlay. Next, samples were treated with 0.25% Trypsin/EDTA for 7 minutes at 37°C and 5% CO_2_. After incubation, trypsin was quenched with cold E8 medium and cell suspensions were centrifuged at 1,200 rpm for 3.5min. Cells were resuspended with FACS buffer (1% BSA in 1X D-PBS) and centrifuged again at 700 rpm for 4 min. Cells were resuspended again with FACS buffer and passed through an EASYstrainer 40 μm filter to ensure single cell suspensions. Resulting suspensions were then analyzed using a BD Biosciences LSR Fortessa flow cytometer and its accompanying BD FACSDiva software. Plots were created from resulting FSC files with FlowJo v10.1.0.0.

### RT-PCR Analysis

Cells cultured by day 3 were processed for FACS similar to flow cytometry. FACS-sorted cells were collected directly into 500uL of Buffer RLT of the RNeasy Micro Kit (QIAGEN, Cat. No. 74004) and RNA isolation was performed according to the manufacturer’s instructions. A NanoDrop 1000 spectrophotometer (ThermoFisher Scientific) was used to determine RNA quality and quantity. Reverse transcription was performed following the Maxima First Strand cDNA Synthesis Kit (ThermoFisher, REF K1671). RT-PCR analysis was performed using TaqMan Universal Master Mix II, with UNG (ThermoFisher, REF 4416710) and TaqMan primers (ThermoFisher, REF 4331182; GAPDH Primer ID: Hs99999905_m1; TFAP2A Primer ID: Hs01029413_m1) on a CFX Connect Real-Time System (Bio-Rad). The 2−ΔCt method was used to quantify relative gene expression with human GAPDH as an internal control. All analyses were performed with at least three biological replicates and three technical replicates.

### Microscopy

All confocal micrographs were acquired the Zeiss LSM 880 confocal laser-scanning microscope and analyzed via Imaris x64 9.2.1 and ImageJ/FIJI version 2.41.

### Publicly Available Datasets

The scRNA-seq data generated in this study is available publicly online and is stored under NCBI GEO Accession GSE288389.

## Supporting information

ReadStats

## Acknowledgements

This work is supported by 2R01HD079546. A.A. is supported by a Graduate Student Fellowship from Eli and Edythe Broad Center of Regenerative Medicine and Stem Cell Research (BSCRC)—Rose Hill Foundation Training Program. We acknowledge the BSCRC Microscopy Core and the Technology Center for Genomics and Bioinformatics (TCGB) Genomics Core. We are grateful to Dr. Ken Yamauchi for his guidance on spinning disk live cell imaging experiments and confocal imaging. We are grateful to Dr. Sissy Wamaitha for her guidance on BMP inhibition experiments and Dr. Varsha Desai for conducting mycoplasma testing. The V.P. laboratory is supported by the Research Foundation-Flanders (FWO grants G0C9320N and G0B4420N to V.P.), KU Leuven Research Fund (C1 grant C14/21/119 to V.P.), and Pandarome project 40007487 (G0I7822N) (funded by the FWO and F.R.S.-FNRS) under the Excellence of Science (EOS) program. T.X.A.P is supported by a FWO PhD fellowship (11N3122N). We thank the staff at the Flemish Supercomputing Center (Vlaams Supercomputer Centrum – VSC) for their support.

## Author contributions

A.A and A.T.C. conceived and initiated the project; A.A designed, performed and quantified most experiments; J.D. and E.J. completed scRNA-seq data analyses and interpretation; A.A. conducted live cell imaging and J.D. helped on image analysis and IF quantification; Y.H. and Q.Y.W. provided the TFAP2A-TdTomato reporter cell line; M.L. conducted flow cytometry and RT-qPCR; T.X.A.P. conducted analyses of multiple datasets with scRNA-seq data under the supervision of V.P.; N.A. and M.S. helped to maintain cell cultures and participated in experiments; N.P.R. provided the FUCS1 reporter cell line; A.A and A.T.C. wrote the manuscript; and A.T.C. supervised the study. All authors edited and approved the manuscript.

---

**Extended Data Fig.1.**
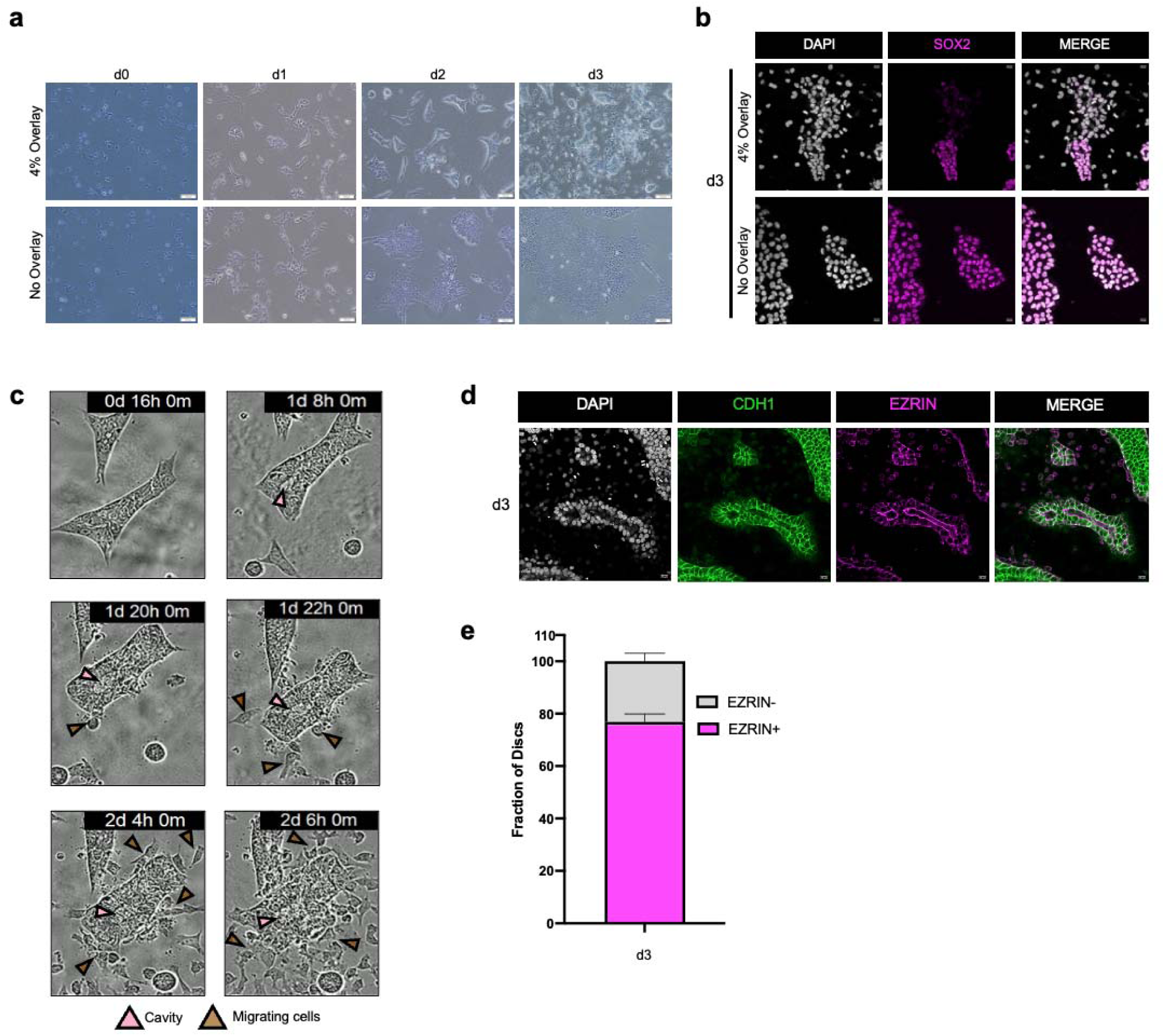
**a,** Phase contrast representative images of hPSCs with and without 4% Geltrex overlay from d0-d3. **b,** Representative images of hPSCs with and without 4% Geltrex overlay at d3. SOX2 (magenta), and DAPI (gray) marks all nuclei present. Scale bar = 20 μm. SOX2 and DAPI had their intensities increased for clarity. **c,** Representative images of phase contrast InCucyte time-lapse video showing GiMO-generated colonies undergo cavity formation and EMT. **d,** Representative images of GiMO-generated colonies at d3. EZRIN (magenta) marks lumen formation and CDH1 (green) marks E-cadherin positive colonies. Scale bar = 20 μm. CDH1, EZRIN, and DAPI had their intensities increased for clarity. **e,** quantification of EZRIN positive colonies generated at d3 GiMO across 2 biological replicates. Bar plot shows mean with SEM.

**Extended Data Fig.2.**
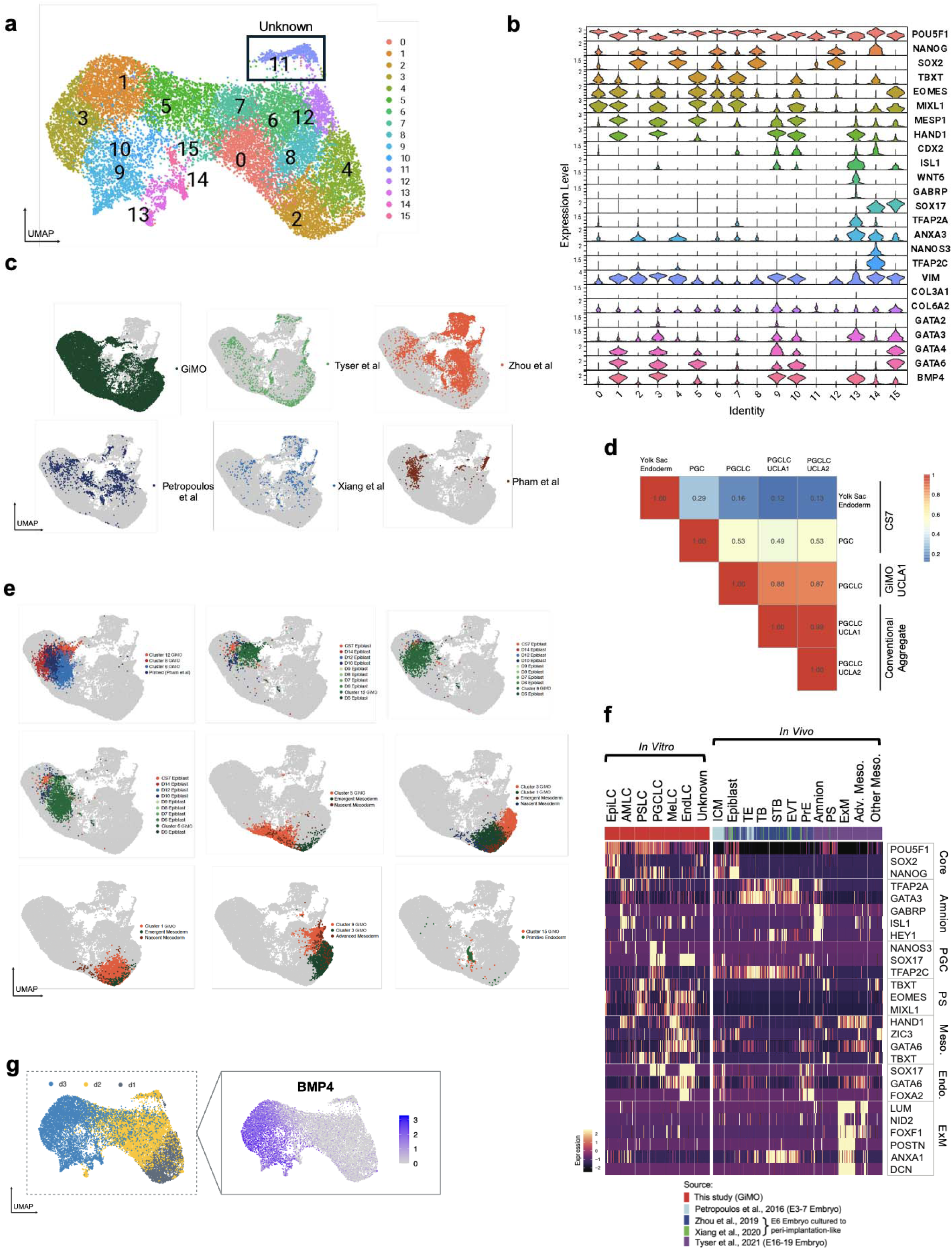
**a,** UMAP plot of scRNA-seq data reveals 16 distinct color-coded clusters, each corresponding cell populations from d1-d3 GiMO differentiation. **b,** Violin plots of marker genes across 16 distinct clusters from d1-d3 GiMO differentiation. **c,** UMAP plot of integrated scRNA-seq datasets from pre-implantation, post-implantation and CS7 human embryo and d1-3 GiMO differentiation. **d,** correlation coefficients among PGCLCs derived from d2-d3 GiMO and PGCs from CS7 human embryo. **e,** UMAP plot of integrated scRNA-seq datasets from pre-implantation, post-implantation and CS7 human embryo and d1-3 GiMO differentiation. **f,** Marker genes expression heatmap integrating GiMO scRNAseq with those from preimplantation, post-implantation and Carnegie Stage 7(CS7) human embryo datasets. **g,** Feature plot showing expression of BMP4 across d1-3 GiMO.

**Extended Data Fig.3.**
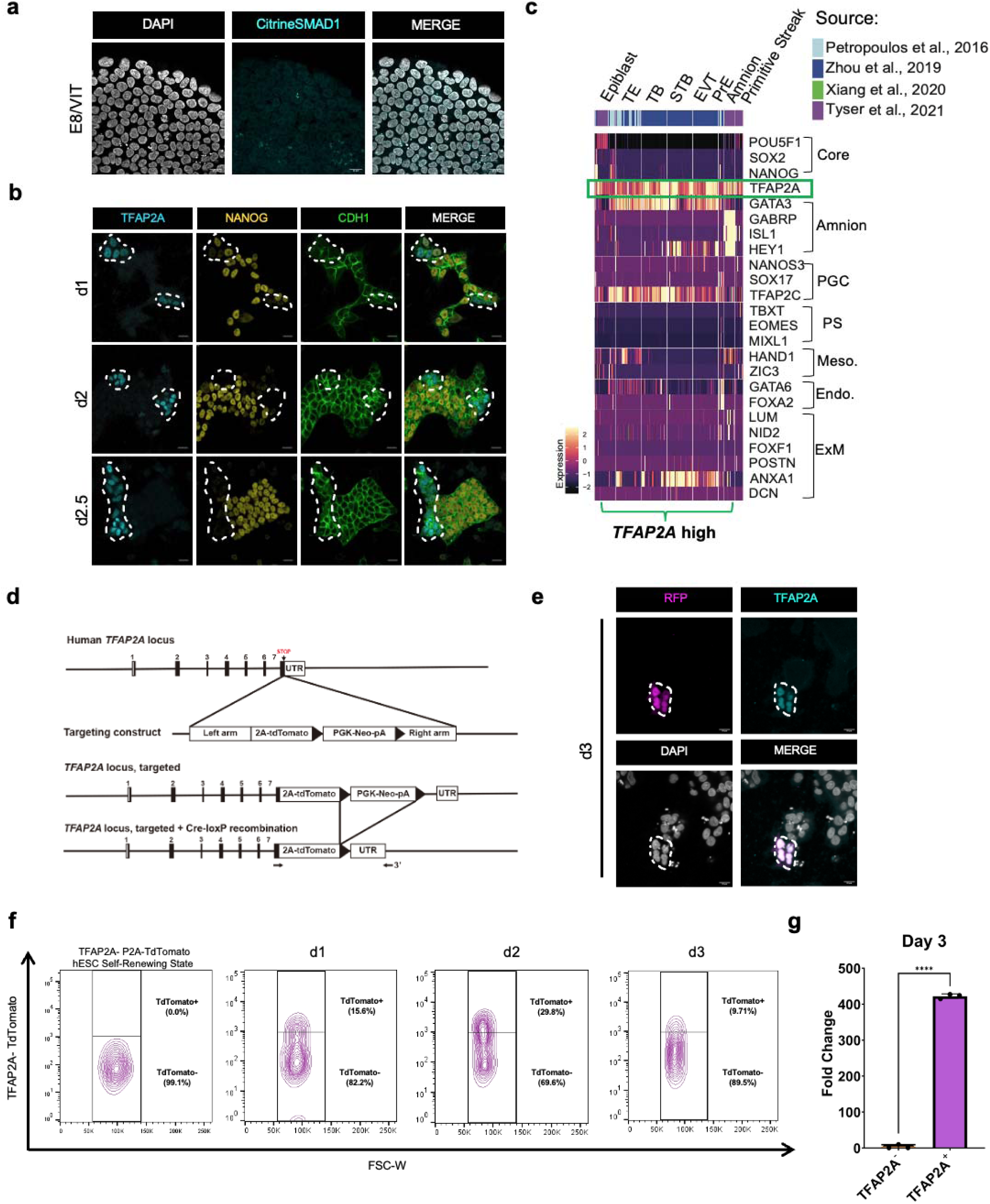
**a,** Representative images of FUCS1 hPSCs under self-renewal state. Cells are maintained on feeder-free Vitronectin coating and E8 media. Cells stained for Citrine-SMAD1 (anti GFP, cyan) and DAPI (gray) marks all nuclei present. Scale bar = 20 μm. Citrine-SMAD1 and DAPI had their intensities increased for clarity. **b,** Representative images of CDH1 (green) colony cells from d1-d2.5 GiMO differentiation showing TFAP2A (cyan) and NANOG (yellow) expression. Scale bar = 50 μm. TFAP2A, NANOG, and CDH1 had their intensities increased for clarity. **c,** Marker genes expression heatmap integrating GiMO scRNAseq with those from preimplantation, post-implantation and Carnegie Stage 7(CS7) human embryo datasets filtering for TFAP2A high expressing cells. **d,** Schematic of inserted vectors for TFAP2A-TdTomato UCLA2 hESC reporter line. Insertion and excision of selection cassettes are shown. **e,** Representative images of RFP (magenta) together with TFAP2A (cyan) at d3 GiMO confirm TFAP2A/RFP overlap. Scale bar = 20 μm. RFP, TFAP2A, and DAPI had their intensities increased for clarity. **f,** flow cytometry analyses of TFAP2A-TdTomato UCLA2 hESC reporter line under self-renewing conditions and d1-d3 GiMO differentiation. Cells were sorted for TdTomato positive and negative expression. **g,** qPCR analysis of TdTomato positive cells FACS sorted at d3 GiMO. Fold change of TFAP2A against hESCs in TdTomato positive versus negative cells. Normalized to GAPDH. N=3 biological replicates. Significance is students T-Test.

**Extended Data Fig.4.**
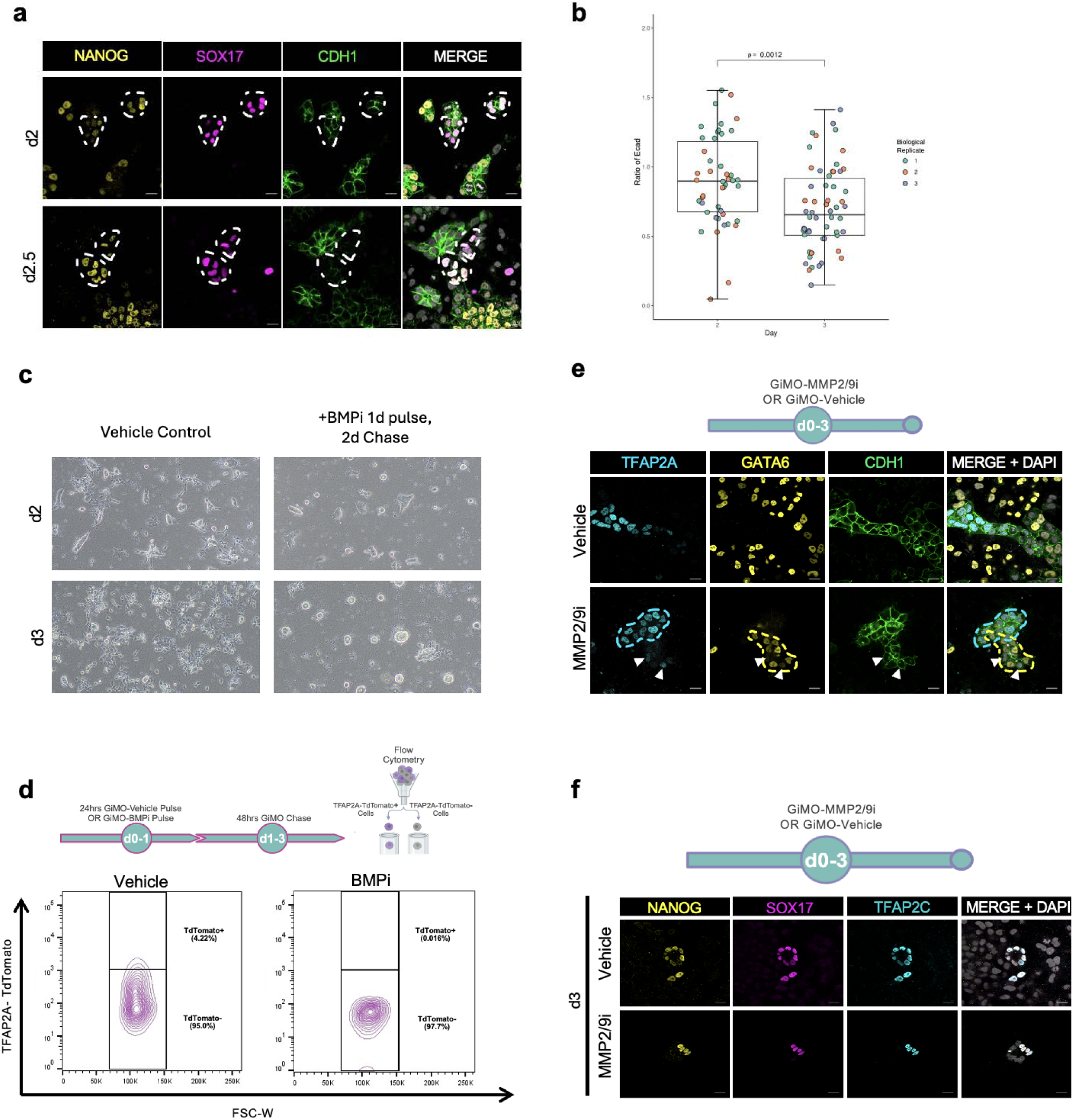
**a,** Representative images of CDH1 (green) colony cells from d2-d2.5 GiMO differentiation showing PGCLC emergence. NANOG (yellow) together with SOX17 (magenta) mark. Scale bar = 50 μm. CDH1, NANOG, and SOX17 had their intensities increased for clarity. **b,** Values in the boxplot are a ratio of mean gray intensities of CDH1 fluorescence in replicate-matched PGCLC-containing and non-PGCLC containing clusters. Boxplot shows whiskers covering full outliers, not 95% confidence interval or SD/SEM. **c,** Phase contrast representative images of Vehicle and BMPi conditions at d2 and d3 GiMO. Scale bars = 200 μm**. d,** Experimental design (top) and flow cytometry analysis (bottom) at d3 GiMO of TFAP2A-TdTomato expressing cells from BMPi pulse-chase (5μm DMH1 concentration) or Vehicle (DMSO equivalent volume). **e,** A schematic of the experimental design (top) and representative images (bottom) of MeLCs and TFAP2A+ cells at d3 GiMO under MMPi (30μm drug concentration) or Vehicle (DMSO equivalent volume). Scale bar = 50 μm. GATA6 (yellow), TFAP2A (cyan), and CDH1 (green) had their intensities increased for clarity. Schematic (top) was made using BioRender.com. **f,** A schematic of the experimental design (top) and representative images (bottom) of PGCLCs at d3 GiMO under MMPi (30μm concentration) or Vehicle (DMSO equivalent volume). Scale bar = 20 μm. NANOG (yellow), TFAP2C (cyan), and SOX17 (magenta) had their intensities increased for clarity.

