## Supplemental Table 1 for "TFAP2A+ embryonic progenitor cells undergo fate diversification to give rise to human amnion, germline, and mesoderm"

| **Antibody (HUMAN)** | **Host Species** | **Dilution** | **Catalog Number** |
| --- | --- | --- | --- |
| Ezrin | Rabbit | 1:250 | Fisher, 50-173-5186 |
| ISL1 | Mouse | 1:200 | DSHB, 39.4D5 |
| E-Cadherin (CDH1) | Mouse | 1:100 | BD, 610182 |
| SOX2 | Rabbit | 1:500 | Abcam, ab97959 |
| GATA6 | Goat | 1:250 | R&D, AF1700 |
|  | Rabbit | 1:250 | Novus, NBP2-55937 |
| NANOG | Rabbit | 1:200 | Abcam, ab109250 |
|  | Goat | 1:250 | R&D, AF1997 |
| TFAP2A (AP2α) | Mouse | 1:100 | Santa Cruz, sc-12726 |
|  | Rabbit | 1:100 | Invitrogen, MA5-14856 |
| TFAP2C (AP2γ) | Mouse | 1:100 | Santa Cruz, sc-12762 |
| SOX17 | Goat | 1:500 | R&D, AF1924 |
| Anti-RFP | Rabbit | 1:200 | Rockland, 200-101-379 |
| Anti-GFP | Chicken | 1:200 | Abcam, ab13970 |
| TBXT (Brachyury) | Goat | 1:500 | R&D, AF2085 |
| LHX1 (LIM1) | Rabbit | 1:350 | Novus, NBP3-21271 |
